# Awareness of the Outcome of Self-Initiated Pointing Actions

**DOI:** 10.1101/234369

**Authors:** Andrei Gorea, Lionel Granjon, Dov Sagi

## Abstract

Are we aware of the outcome of our actions? The participants pointed rapidly at a screen location marked by a transient visual target (T), with and without seeing their hand, and were asked to estimate (E) their landing location (L) using the same finger but without time constraints. We found that L and E are systematically and idiosyncratically shifted away from their corresponding targets (T, L), suggesting unawareness. Moreover, E was biased away from L, toward T (21% and 37%, with and without visual feedback), in line with a putative Bayesian account of the results, assuming a strong prior in the absence of vision. However, L (the assumed prior) and E (the assumed posterior) precisions were practically identical, arguing against such an account of the results. Instead, the results are well accounted for by a simple model positing that the participants’ E is set to the planned rather than the actual L. When asked to estimate their landing location, participants appeared to reenact their original motor plan.

## INTRODUCTION

How well do we actually ‘know’ the outcome of our actions? To the best of our knowledge, this question was asked for the first time by Goodale, Pelisson, & Prablanc (1986) and Fourneret & Jeannerod (1998) and was answered negatively: participants requested to draw a straight line on a tablet were not aware of or they greatly underestimated the unbeknown to them perturbation of their motor outcome. Although many studies have demonstrated thereafter the existence of a corollary discharge (Ariff, Donchin, Nanayakkara, & Shadmehr, 2002; Miall, Christensen, Cain, & Stanley, 2007; Sommer & Wurtz, 2008; Wagner & Smith, 2008), which, within the framework of forward models (Wolpert & Ghahramani, 2000), should allow for such introspective knowledge, only recently has this issue been addressed again but with mitigated results. van Dam & Ernst (2013) showed that participants that had performed a one-dimensional hand-pointing task were beyond chance when they were later asked to decide whether their landing location had been left or right of the target. The same study, however, showed that participants had no knowledge of the size of their pointing error because they were unable to correct their landing location. However, none of these studies focused on the participants’ putative introspective bias. This issue has only been recently addressed using a paradigm involving only the participants’ key press (hence, not involving a proper motor plan) in order to align a moving dot with a static reference (see below; Wolpe, Wolpert, & Rowe, 2014). That study revealed that the participants believed that they were better than they actually were (i.e., they minimized their alignment error), a bias referred to by the authors as an “exaggerated expectation of success”. In a similar vein, many studies have shown that humans tend to overestimate their intellectual and social performances (Dunning, Johnson, Ehrlinger, & Kruger, 2003), let their perceptual decisions be biased by induced mental sets (Sekuler & Ball, 1977) or by placebo manipulations (Sterzer, Frith, & Petrovic, 2010) and generally are prone to let their cognitive and sensory decisions be biased by their expectations (Seriès & Seitz, 2013), by additional congruent or incongruent sensory cues(Ernst & Banks, 2002), or by their feelings of agency (Desantis, Waszak, & Gorea, 2016; Hughes, Desantis, & Waszak, 2013).

In contrast with such systematic and shared sensory and cognitive biases, it has been shown that in the presence of a target but in the absence of visual feedback during a pointing task, participants display idiosyncratic pointing biases, presumably resulting from systematic errors in their sensorimotor transformation from the visual representation of the target location to the kinematic representation of their arm movement (Chang & Snyder, 2010; Sober & Sabes, 2003; Soechting & Flanders, 1989a, 1989b). Systematic idiosyncratic biases have also been reported for directional preferences in eye movements, motion perception (Schütz, 2014) and bistable perceptions (Wexler, Duyck, & Mamassian, 2015). Whether these biases are actually involved in participants’ estimation of the outcome of their own actions remains unknown.

The question concerning the introspective knowledge of one’s action outcomes and of its putative bias is investigated here in the context of a manual pointing task. The answer is provided by the characteristics of the distribution of participants’ estimates of their own pointing locations. Although theoretical accounts of introspective knowledge (van Dam & Ernst, 2013) and of many kinds of perceptual and motor biases have appealed to the Bayesian interpretative framework (Trommershäuser, Maloney, & Landy, 2008; Wei & Stocker, 2015, 2016; Weiss, Simoncelli, & Adelson, 2002; Wolpert, 2007; Wolpert & Ghahramani, 2000; Wolpert & Landy, 2012), here we show that the Bayesian approach is incompatible with the present results. We propose an alternative approach exclusively based on the general forward model framework (Shadmehr, Smith, & Krakauer, 2010). Note that our question (as well as that asked in previous related studies (Fourneret & Jeannerod, 1998; van Dam & Ernst, 2013; Wolpe et al., 2014) and the experimental design presently used to address it do not pertain to how accurately people can reproduce their (ballistic) motor actions but instead pertains to how well they *know* the *outcomes* of such actions. For example, had the motor action been a juggler’s hand motion to receive the prop, our question would not have been how accurately the juggler could reproduce that precise movement (she reproduces it with sufficient precision countlessly) but how well she could specify the location of her hand when about to grasp the prop in the absence of visibility of the prop. This distinction is critical insofar as the reproduction of a motor action under the same conditions as the original requires the recovery of the whole motor sequence from memory, whereas localizing the hand involves knowledge of the outcome of that sequence.

Consider a single pointing and estimation trial where the target is displayed at a random location (in a frontal plane) and disappears before the hand starts moving. This results in three locations on the plane, those of the Target, T, of the finger’s landing, L, and of participant’s estimated landing location, E (Figure 1). If participants estimate L independently of T, then over trials, L and E should behave as *independent* random variables and could possibly display independently biased means (constant errors relative to their corresponding T and L locations, respectively). In addition to such putative constant errors, a trial-by-trial correlation between these variables (i.e., between vectors 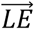 and 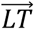) would imply that E estimates are affected by expectations related to the goal of L, which is biased by the target T. Therefore, in the context of the task described, with regard to the question of how well we know the outcome of our actions, the answer involves three E-related measurements: its precision, i.e., 1/standard-deviation of our *meta-* knowledge, the *consistent error,* i.e., the accuracy of our meta-knowledge, and the correlation between 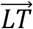 and 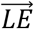, an additional meta-knowledge measure of accuracy reflecting the extent to which our knowledge is *biased* by the goal of our actions.

**Figure 1.**
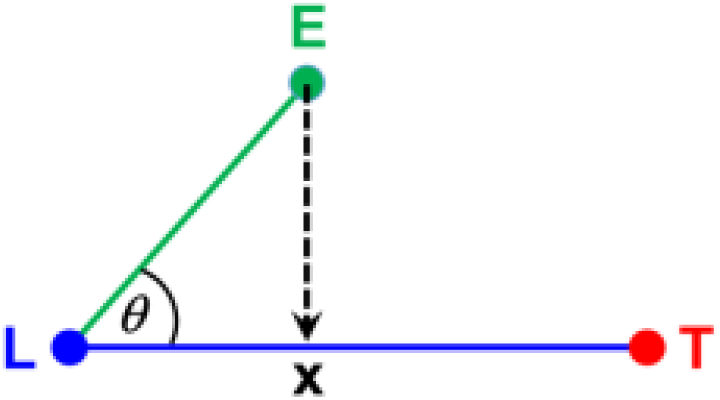
*The Target (T), Landing (L), and Estimated landing (E) triplet with the projection of E on the LT̅ vector, x. For E being independent of T, x is expected to have an isotropic distribution around L, with <x>=0.*

In principle, a putative E bias (i.e., toward the goal of our action, T) could be accounted for within a Bayesian decision framework where information related to the task (i.e., the prior) is combined with the actual measurement, yielding a higher precision estimate. In a previous study (where a mouse-click was used to stop a drifting dot in order to vertically align it with a marker, as well as to subsequently estimate the location of the ‘stop’; Wolpe et al., 2014), the Bayesian framework was successfully fitted to the data. Our approach of analyzing and modeling the present data allowed us to critically evaluate this theoretical framework, in particular, with respect to one’s evaluation of self-initiated action. Our theoretical approach is based on an oversimplified *internal model.* It posits that, under the experimental conditions tested here (a speeded up pointing task with and without visual feedback), the participants access the location P of their *planned* movement, around which they land, so that instead of estimating their actual landing, they estimate P (with a given precision). Based on this theoretical account, both L and E gravitate around P with independent noises and biases (Soechting & Flanders, 1989a, 1989b). Importantly, we suggest that participants are not aware of their L and E biases.

## MATERIAL AND METHODS

All experiments were performed in accordance with relevant institute guidelines and regulations. The protocol of the experiments was approved by the Paris Descartes University Ethics Committee for Noninvasive Research (CERES). Informed consent was obtained from all participants.

### Stimuli

The target stimuli and a 3.2 deg (72 px / 19 mm) white (90 cd/m^2^) cross designating the finger starting position were presented in a dimly lit room on a Cintiq 24HD touch tablet (active area: 518×322 mm; 1920×1200 px; 75 Hz raster). A chin rest was used to control the viewing distance so that the participants’ eyes were 35 cm in front of the vertically mounted touch screen. The computer keyboard was placed in front of the touch screen and slightly eccentrically with respect to it to facilitate the participant’s pressing of the spacebar (see Figure 2). The start-cross was placed centrally on the touch screen and 4.9 cm (180 px) from the bottom of the Cintiq’s active area. The target stimuli were either 1 deg (24 px / 6 mm) radius, a white disk (90 cd/m_2_), or a low spatial frequency luminance noise pattern windowed by a 2D Gaussian envelope with a SD of 4.2 deg (100 px / 27 mm) (average luminance of 60 cd/m_2_). The center of these targets could appear anywhere within one of three rectangular arrays (1.8×5deg / 41×113 px / 11×30 mm) with their centers placed on a virtual half-circle of radius 13 cm/482 px/21.3 deg centered on the finger’s starting location (see Fig. 2). The mid-rectangular array was placed straight ahead (i.e., at 90° with respect to the bottom edge of the tablet) with the other two location centers at ±45° on the virtual circle. The center of each rectangle was offset over trials by a random jitter uniformly distributed horizontally and vertically at a range of ±1.6 cm/59 px. The experimental software controlling the target display and the response recordings was implemented in MATLAB using the Psychophysics toolbox (Brainard, 1997; Pelli, 1997).

**Figure 2.**
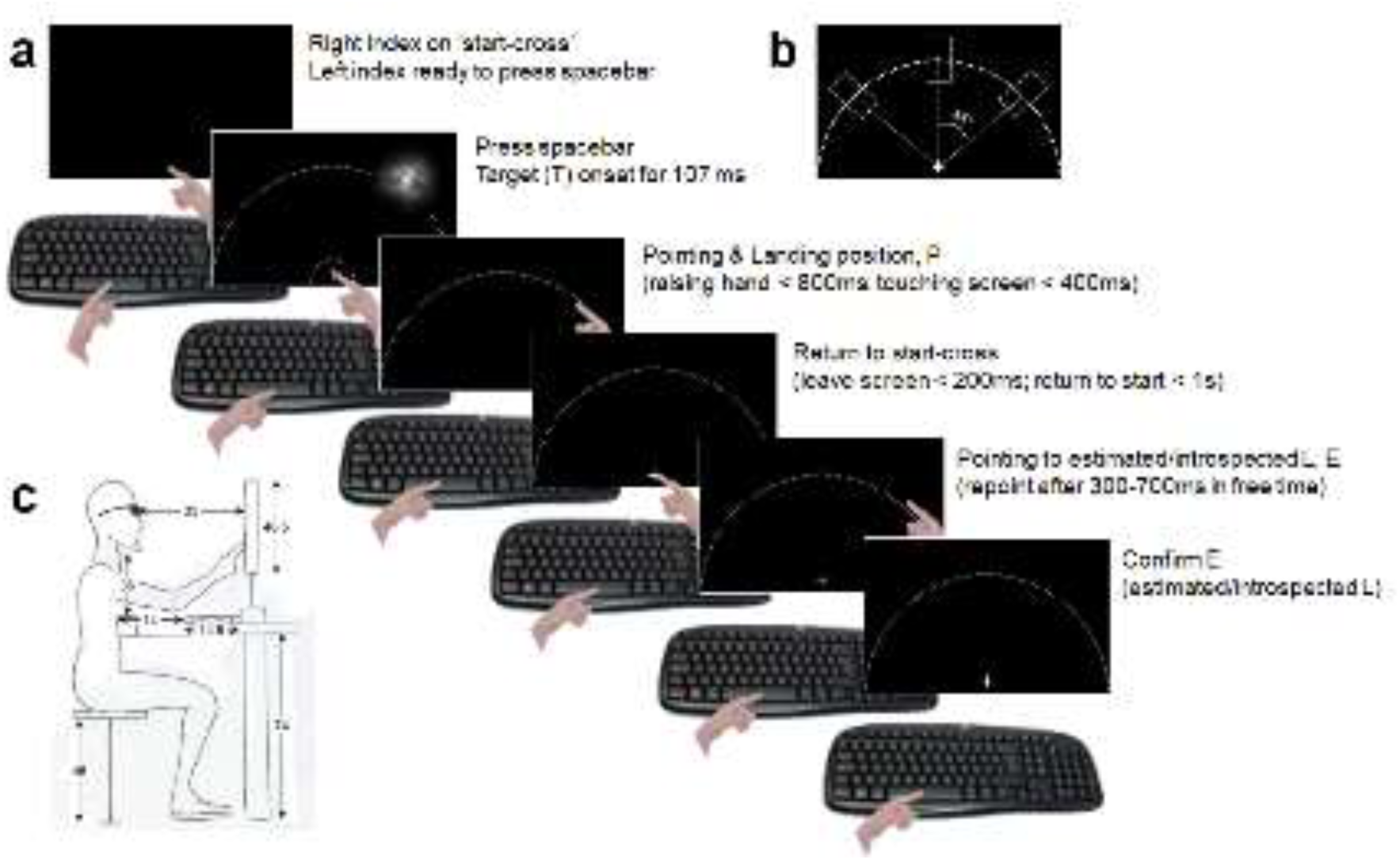
*Experimental setup. **(a)** One trial showing the extended target (P-conditions). **(b)** The three arrays specifying the possible target locations. **(c)** A participant with shutter PLATO glasses facing the screen with dimensions given in cm. (All parts of this figure were drawn – the hand – or designed by one of the authors – AG – and by Esther Bernager – the diagram in c). The photograph of the keyboard was also taken by AG.)*

All the spatial parameters above and the temporal parameters below were set, based on preliminary trials run by two of the authors to minimize fatigue and facilitate as much as possible the pointing movements while still preserving a reflexive motor behavior.

### Participants

Fourteen participants (7 female, 1 left-handed) between 21 and 25 years of age took part in all the experiments. They had no history of neurological disorders and had normal or corrected-to-normal vision.

### Procedure

Participants had their head posed on a chin rest and were seated so that their eyes were 35 cm from the tablet screen and the elbow of their dominant hand resting on the table. In all experiments one trial was divided into two phases: the target-reaching phase and the (introspective) estimation of the landing location phase. The target-reaching phase started with the participant having the index finger of the dominant hand (only one was left-handed) on the starting cross, and with the other hand ready to press the spacebar. With a delay of 500-900 ms, the spacebar press triggered the appearance of the target stimulus at one of three possible locations. The target was extinguished after 107 ms. Participants were instructed to point at the target location with their index finger as fast as possible while raising their elbow from the resting position. Raising their index from the start-cross with a delay longer than 800 ms or touching the screen more than 400 ms after having raised their index finger from the start-cross was considered as a failed trial, which was replaced later in the sequence. Participants had to quit their landing position within less than 200 ms and reposition their index within a radius of 4.8 deg/108 px about the center of the start-cross within less than 1000 ms. Noncompliance with either of these two durations was considered as a failed trial, which was also replaced later in the sequence. The landing position estimation phase started once the participant had repositioned her finger on the start-cross: 300 to 700 ms later, a brief tone prompted the participant to quit the start-cross and to point at the screen location that she thought was her landing location. To validate their choice, the participants pressed any keyboard key except the spacebar with their free hand. The time allowed for the estimation phase was unlimited but participants took on average 0.97 ± 0.26 sec.

Participants were run under four experimental conditions, namely, with the spatially extended targets (referred to in the text as P-) and with the dot-targets (referred to as P+) and with full (V+) and with limited vision (V-). The use of P-conditions was primarily intended to explore the dependence of the introspective estimation bias on the spatial precision of the target. Participants were instructed to point at any location within the spatially extended targets and to be as precise as possible when pointing at the dot-target. The limited vision (V-) was applied only to the target-reaching phase, where vision was totally obstructed 0 ms after the participant’s index finger left the start-cross and was restored as soon as the pointing finger touched back the start location. Visual obstruction was obtained by means of PLATO Visual Occlusion Spectacles(Milgram, 1987) (Translucent Technology, Inc.), whose special characteristic is that they preserve the average ambient luminance.

Each of the four experimental conditions (P+V+, P+V-, P-V+, and P-V-) was blocked within one experimental session (with the three target-array locations randomized within the session) and repeated four times in an order randomized across the participants. Each session was split into two blocks of trials: a training block and an experimental block. Each initial training block for each experimental condition consisted of 30 valid trials for each of the three target arrays (i.e., 90 training trials/condition). About 8% of the trials were rejected based on the constraints mentioned above and were repeated in random order to achieve the 30 valid trials. Each of the remaining three training blocks per condition consisted of 10 valid trials per target location (90 trials/condition excluding less than 5% rejected trials). Hence, in total there were 180 valid training trials per condition (totaling 720 training trials). In all cases one experimental session (i.e., each repeated condition) consisted of 35 valid trials for each of the three target arrays repeated four times (i.e., 420 trials/condition), hence, totaling 1680 experimental trials. The entire experiment took about 4 and a half hours per participant. Participants ran only four (out of 16) sessions per day and the whole experiment was completed in no more than 2 weeks. In the final analysis, trials with estimation times longer than ±3.5 SD of the mean (per participant/condition) were discarded (1.5% of total trials). The rejected trials added only 0.51% to the SD estimation (relative to L).

## RESULTS

### Raw data

As expected, landing and estimation locations were elliptical and elongated along the hand-finger motion axis(Sober & Sabes, 2005; Van Beers, Sittig, & Denier Van Der Gon, 1998) with average orientations of 149.5(135), 93.5(90), and 35.1(45) degrees for the left, center, and right rectangular target arrays (in between parentheses are the actual orientations of the target arrays). Their average major-to-minor Landing/Estimation ratios were 1.44/1.45, 1.74/1.72, and 1.33/1.33 for left, center, and right target locations, respectively. Their compound standard deviations (i.e., the square-root of the summed squared SDs along the major and minor axes) are given in the section *Landing and Estimation precision.*

The participants performed the speeded up pointing task with a mean reaction time of 0.544 sec, with the finger movement starting at 0.31±0.03 sec (mean and SD across the 13 participants, 4 conditions, and 3 locations) after the target onset, and touching the screen 0.24±0.04 sec later. The average estimation time was 0.97±0.26 sec. Pointing and estimation performances are given below in mm with 1mm = 0.16 deg.

### Consistent L and E errors

Figure 3A exemplifies for two participants what we mean by L and E *consistent errors.* The black rectangles (not presented in the experiments) circumscribe the areas within which a target (black dots, corresponding to the centers of the presented targets) could be displayed (Left T-arrays in the illustration). The blue and red rectangles reproduce the black ones with their centers positioned at the corresponding centers of the L and E trial clusters for conditions P+V+ and P+V-(rows) and for two participants (columns). The global displacements of the blue and red rectangles with respect to the black and blue ones, respectively, are what we refer to as L (with respect to T) and E (with respect to L) consistent errors. These global shifts are idiosyncratic. Figure 3B displays the centers of the three clusters (T, L, and E, colored correspondingly) for the 4 conditions, the three locations, and 13 participants, with each pair of blue and green circles (linked by straight lines) representing one participant. Note that the T locations vary slightly across participants and conditions, since the jitter of the target within the rectangular T-arrays adds randomness to the trial-by-trial target display. Each blue and red circle center marks a location averaged over all trials (per participant, condition, and location) with the standard error of the mean E locations smaller than half of the circle’s radius. As such, Fig. 3B clearly shows highly significant L (with respect to T) and E (with respect to L) consistent errors. If the participants had not displayed any such consistent errors, their mean L and E should have not differed significantly from T and L, respectively. However, this was clearly *not* the case, with consistent errors being, on average, 16.7 times the SEM (p < 0.01 for 309 out of 312 measured means).

**Figure 3.**
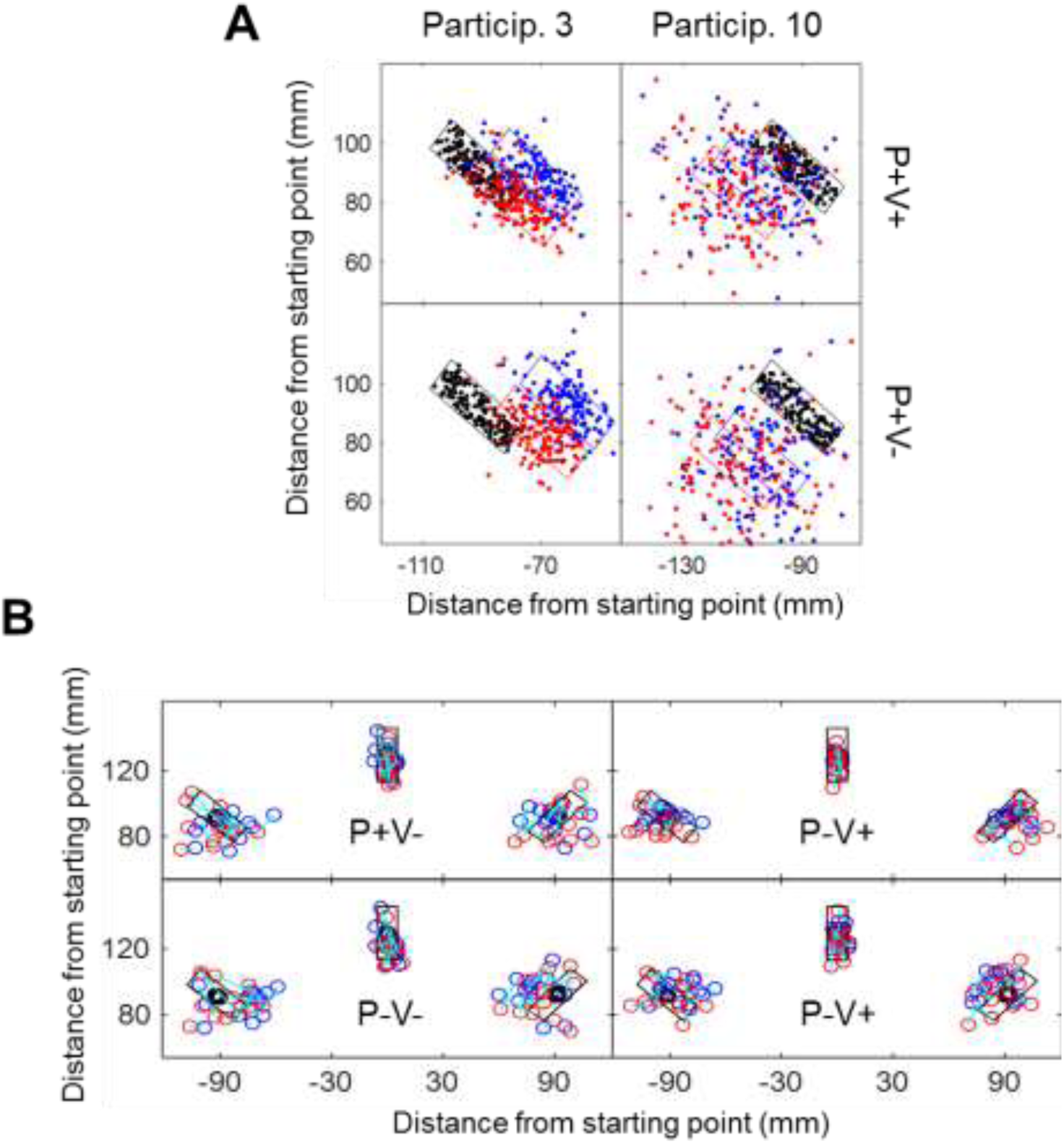
**A.** *Target (T), Landing (L), and Estimated L (E) locations (respectively, black, blue, and red dots and rectangles) for two participants (columns), for conditions P+V+ and P+V-(rows) and for the left-side T-arrays. Black rectangles delimit the T-arrays, with the blue and red rectangles reproducing the black one but with their centers shifted according to the mean L and E locations. **B.** Mean T, L, and E locations (black, blue, and red circles) computed for each of the 13 participants for the 4 experimental conditions (different panels) and for the 3 T-arrays, with the latter shown as black rectangles. Linked blue and red circles are for the same participant. The diameters of the circles are at least 4 times the standard error of their represented mean location. X and Y coordinates are in mm relative to the movement’s starting point.*

To quantify the similarity of the participants’ consistent errors across the four experimental conditions, we estimated the across-participants variance shared by all conditions, using principal component analysis (PCA, using Matlab©). For each of the four conditions there were 39 measurements (13 participants × 3 locations), forming 4 data series of 39 measurements each. Each series was z-scored before PCA was applied. The analysis shows that for the L consistent error (relative to T), the first principal component explains 76% of the data variance, and for the E consistent error (relative to L) the first component explains 89% of the data variance. We performed two control tests; in test 1 the participants were randomly shuffled within each condition (keeping the three locations aligned, though with different participants’ results); in test 2 the results were randomly shuffled within each condition. For test 1, the first component accounted for 39 ± 2.7% (mean ± SD computed over 10000 repetitions) of the L variance, and for 38 ± 3.1% of the E variance. For test 2, the first component accounted for 35 ± 2.4% of the L variance, and for 35 ± 2.4% of the E variance. The outcomes of both test 1 and test 2 are comparable to those obtained with an uncorrelated normally distributed data series of the same size, namely, 34 ± 3.1%. When combining the L and E data series (totaling 8 series of 39 measurements each), the first principal component accounts for only 54% of the data, and the second contributes an additional 32% of the variance, suggesting some differences between L and E consistent errors. Here, test 1 and test 2 yield the first components accounting for 28 ± 1.2% and 22 ± 1.4% of the data variance, respectively, comparable with the 22 ± 1.8% yielded by random sequences. These analyses clearly show that consistent errors are participant dependent and are very similar across experimental conditions.

### Landing and estimation precision

Figure 4 displays the standard deviations (SD = 1/precision) of L with respect to T (σ_LT_) and of E with respect to L σ_EL_ and to T (σ_ET_) averaged over the 13 participants. In the presence of vision (V+), both the L and E precisions are higher (the SDs are smaller) than in its absence by an average of 17% and 22%, respectively (both significant at p <<.0001; one-tailed paired t-tests). For L, this is clearly a result of participants being able to better monitor their pointing behavior when their vision of the moving hand is available despite the rapidity of their arm movement (Desmurget & Grafton, 2000; Gordon & Ghez, 1987; Paillard, 1996; Prablanc & Martin, 1992; Sabes, 2000; Sarlegna et al., 2003, 2004, Saunders & Knill, 2003, 2004). For E (relative to L), for which vision is always available, the improved precision may result from participants being able to visually localize their landing finger (i.e., L). The precision of the target (P+ vs. P-) improves L and E precisions by 15% and 5% (p = 0.006 and 0.008, respectively, one-tail, pairwise t-tests). Given that the average pointing SDs to 1 deg (P+ condition) and to 4.2 deg (P-condition) radius targets were respectively, 0.41 and 0.48 deg (i.e., a 15% precision gain for P+), one can speculate that in the latter case (P-) participants aimed at an area of about 1.2 deg radius within the extended 4.2 deg radius P-area. The E precision is not expected to depend on the target size unless, somehow, the participants try to match their estimation precision to their landing one (implying that they have some knowledge of the latter). The noise parameters obtained using a *Simplified Internal Model* (presented below) allowed us to better understand the hidden processes accounting for the observed results.

**Figure 4.**
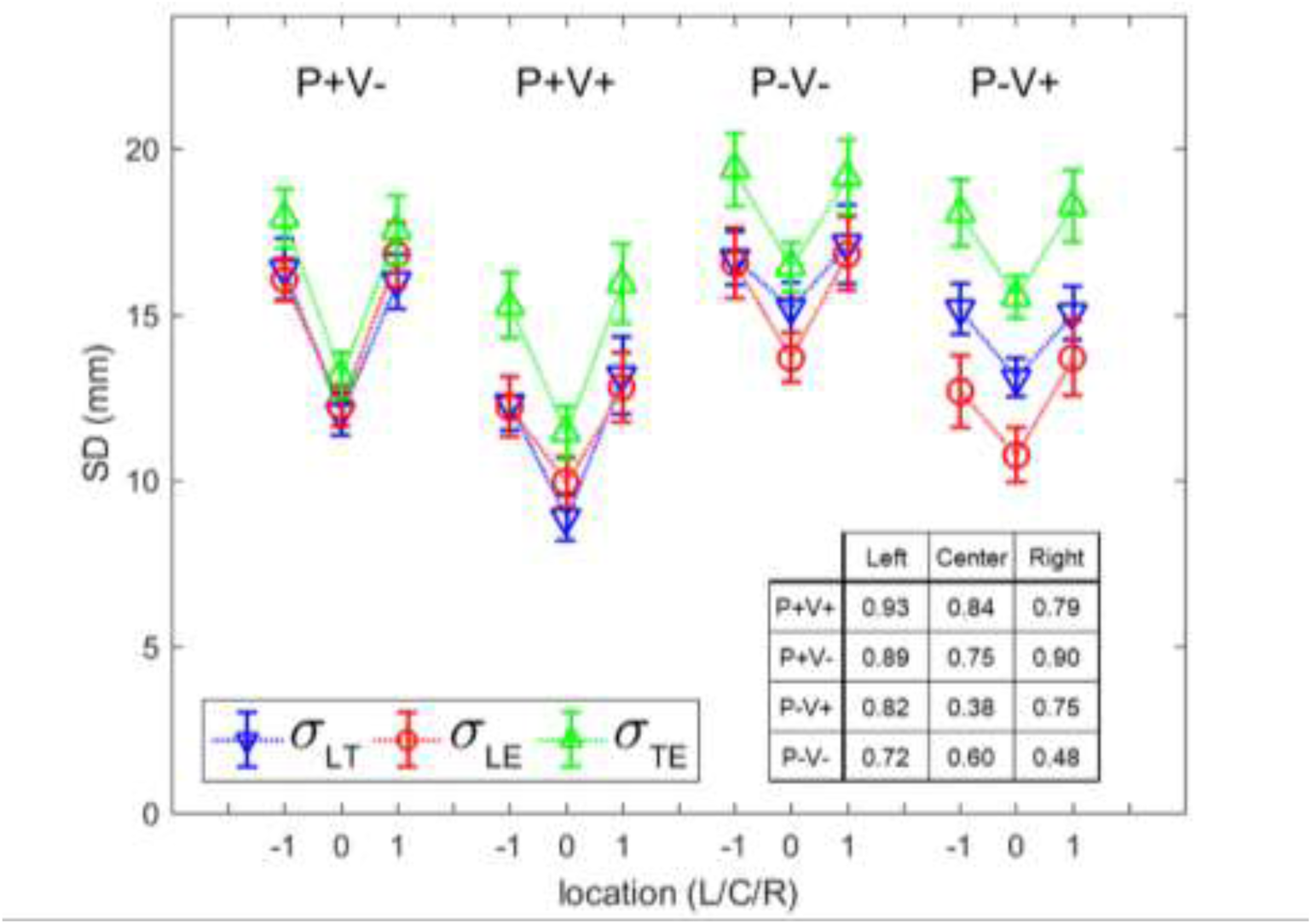
*Standard deviations (i.e., the inverse of precision) in mm of L relative to T (σ_LT_; inverted triangles) and of E relative to L and to T (σ_LE_, σ_TE_; circles and upright triangles) averaged over the 13 participants for each of the four experimental conditions and for each of the 3 T-arrays (marked −1, 0, and 1 for Left, Center, and Right T-arrays; L/C/R). The inset shows the across-participants’ correlations between σ_LT_ and σ_LE_ (all but two, r = .38 and r = .48, are significant at least at a p<0.05 level). Error bars are ±1SEM (N=13).*

An interesting feature of the data displayed in Fig. 4 is the practical equivalence of σ_LT_ and σ_LE_, with the latter only slightly smaller than the former, but inconsistently so across conditions. Within a Bayesian decision-making framework, the E distribution (around L) is taken to be the posterior and must be narrower than the prior distribution instantiated by the L distribution (about T; including visual uncertainty in the presence of sensory information). This is not the case for the average data presented in Fig. 4 for all three locations and for two (P+V+ and P-V-) out of the four conditions, a result incompatible with a straightforward Bayesian approach. When considering each participant individually, there are 67 such Bayesian incompatible cases out of 156 (42.9%). This leads to four possible conclusions, namely that (i) there are distinct Bayesian compatible and Bayesian incompatible populations, (ii) the incompatible cases reflect a measurement error and that the prior and posterior distributions have in fact equal precisions, (iii) the estimation error includes a source of error not taken into account by a straightforward Bayesian approach, or, finally, (iv) overall, the Bayesian framework cannot account for the present data. A more detailed inspection of the data rejects the first possibility because each of the 13 participants shows at least 4 cases (out of 12) with posterior precisions smaller than their prior precisions. The second possibility implies an infinitely large likelihood variance and hence, a posterior centered on the prior (i.e., the target) rather than on the landing location. The data do not support this prediction either, as discussed below in the section *Estimation bias.* Finally, the last two possibilities will be addressed in some detail in the Discussion section.

A final observation about the L and E precisions is that they are significantly correlated across participants for each of the four experimental conditions and for each of the three T-arrays (with 2 exceptions, i.e., r = 0.38 and 0.48 - out of 12 conditions; see the inset in Fig. 4) at p levels of at least 0.05 (computed over the 13 participants). In agreement with previous reports (Wolpe et al., 2014), these correlations show that the precision of humans’ actions is paired with their precision in estimating the outcome of those actions.

### Estimation bias

The estimation bias refers to the extent to which the location of E depends on the location of T. In principle, L (with respect to T) and E (with respect to L) should be independent random variables. Their interaction should be manifested by a significant 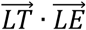 correlation across trials. A positive correlation implies that E is in the direction of T relative to L. These correlations were computed over trials of each participant and for each of the 4 conditions and 3 T-arrays (N = 140). Their means across participants are shown as circles in Figure 5. These correlations are all positive and highly significant (p = 0.00013, sign test separately performed for each correlation value), showing that, on average, E is shifted toward T (mean 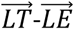 angle - indicating an orthogonal bias relative to 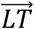 - not significantly different from 0), thereby supporting the notion of an “exaggerated expectation of success”, i.e., the participants’ belief of having pointed more accurately (closer to the target) than in reality(Wolpe et al., 2014). 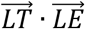 correlations do not depend on the participants’ precision, showing only weak correlations with the related standard deviations (σ_LE_ and σ_LT_) with r values ranging between 0.23 and 0.3 (computed over 13 participants’ × 3 locations per condition, i.e., N = 39; all p > 0.05).

**Figure 5.**
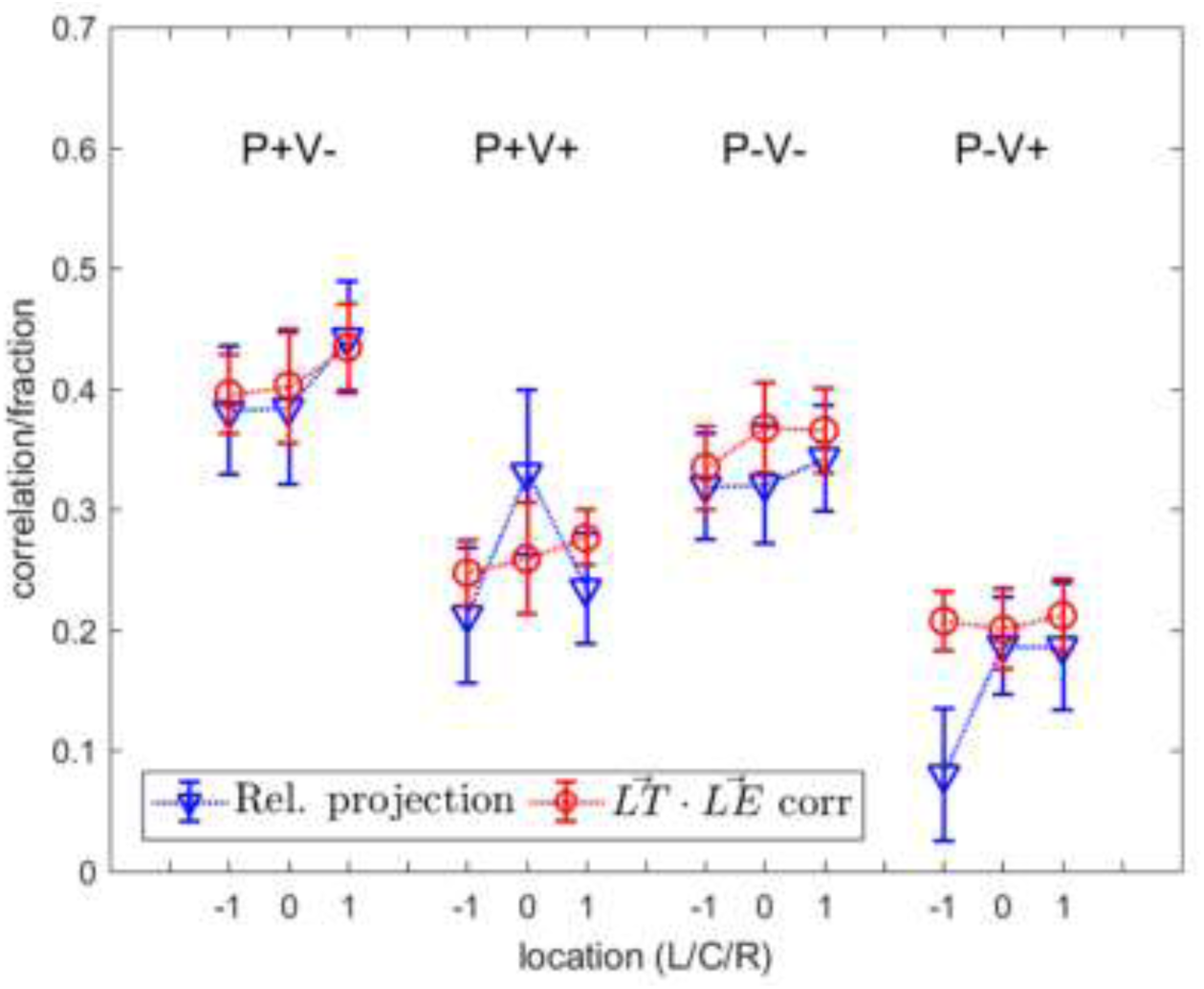
*Mean relative trial-by-trial projections of the 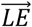 vector on the 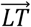 vector (triangles) and 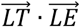 correlations across trials (circles) averaged over the 13 participants for each condition (labels) and location (with −1,0,1 standing for Left, Center, and Right T-arrays; L/C/R). Error bars are ±1SEM (N = 13). Note the close similarity of the correlation and projection values, though their computation methods differ (mainly using the normalization method; thus, the similarity is expected for similar σ_LE_ and σ_LT_ values).*

The 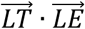 correlation across trials is blind to the consistent errors (i.e., the correlation function ignores the mean). An alternative measure of the E bias including the consistent error is given by the trial-by-trial projection of the 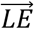 vector on the 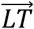 vector, given as a proportion of the 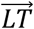 vector (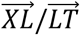 in Fig. 1). In the absence of any bias, the average relative projections should be zero. If participants systematically confuse their landing location with the target location, the relative projection should be 1. In the absence of consistent errors, the Bayesian operator predicts a relative projection of 1 when L precision equals E precision (i.e., the prior equals the posterior) as is actually assessed (see the section *Landing and estimation precision).* The trial-by-trial relative projections were averaged for each participant, condition, and T-array and then averaged over participants for each condition and for each T-array, before and after subtracting the consistent errors. Averaged over all participants, conditions, and T-arrays (13 × 4 × 3 estimations), the relative projections were 0.25 and 0.29, respectively before and after subtracting the consistent errors but with a much smaller dispersion for the latter case (σ = 0.40 vs. 0.19). The relative projections in the presence of vision show significantly less bias (0.37 before and 0.21 after subtraction) than in its absence (0.43 before and 0.37 after subtraction; both two-tailed paired t-tests with p <<.0001, N=78). The relative projections under the imprecise target condition (P-) were 49% larger (p=0.001; paired t-test) relative to the precise target condition (P+). These relative projections are shown as triangles in Fig. 5. They are all significantly larger than 0 and smaller than 1 at p levels < 0.01 or less (N=13 participants, sign tests), meaning that participants’ E systematically deviates from L toward T but does not coincide with T. Note that the 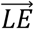 projection on 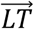 ignores the bias in the orthogonal direction (i.e., orthogonal to the 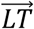 vector). The projection of 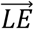 on this axis is not appreciably different from 0 and is also much less dispersed after (σ = .27) than before (σ =.46) subtracting the consistent errors.

### A simple Internal Model

The correlations between 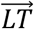 and 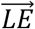 and the relative projections of E on 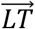 shown above (Fig. 5) rule out the intuition according to which the estimation is unbiased, that is, it is performed around the landing location independently of the target location. This implies that pointing and estimation share a common source of variance. The present data appear to be compatible with the Bayesian view of a prior distribution centered on the target. However, given the approximate equality of the landing and estimation dispersions, the Bayesian account predicts a complete reliance on the prior distribution and hence, an estimated landing location equivalent to the target location. However, our data indicate otherwise. We suggest that the participants’ landing location, L, is distributed around the *planned* landing point, P, which can be viewed as a noisy internal representation of T (distributed around T with a given planning error, Op), and that participants believe that L is at the planned landing location, P. Hence, estimations (E) are distributed around P. The quantitative implementation of this scenario successfully describes our pointing and estimation results related to precision and bias. The model ignores the idiosyncratic consistent errors in pointing (L) and estimation (E) reported in the Results section (Fig. 3), since these are mostly participant dependent, are assumed to be the outcome of motor biases not available to the participants’ introspection, and do not affect the measured precision and bias.

The model (Figure 6) involves three independent random variables: P, L, and E, with P distributed around the target (T), with SD = σ_P_ and with both L and E distributed around P with SD σ_L_ and σ_E_ Given the above description of events, the variances in the data can be written as:

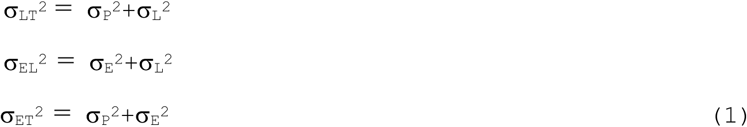

**Figure 6.**
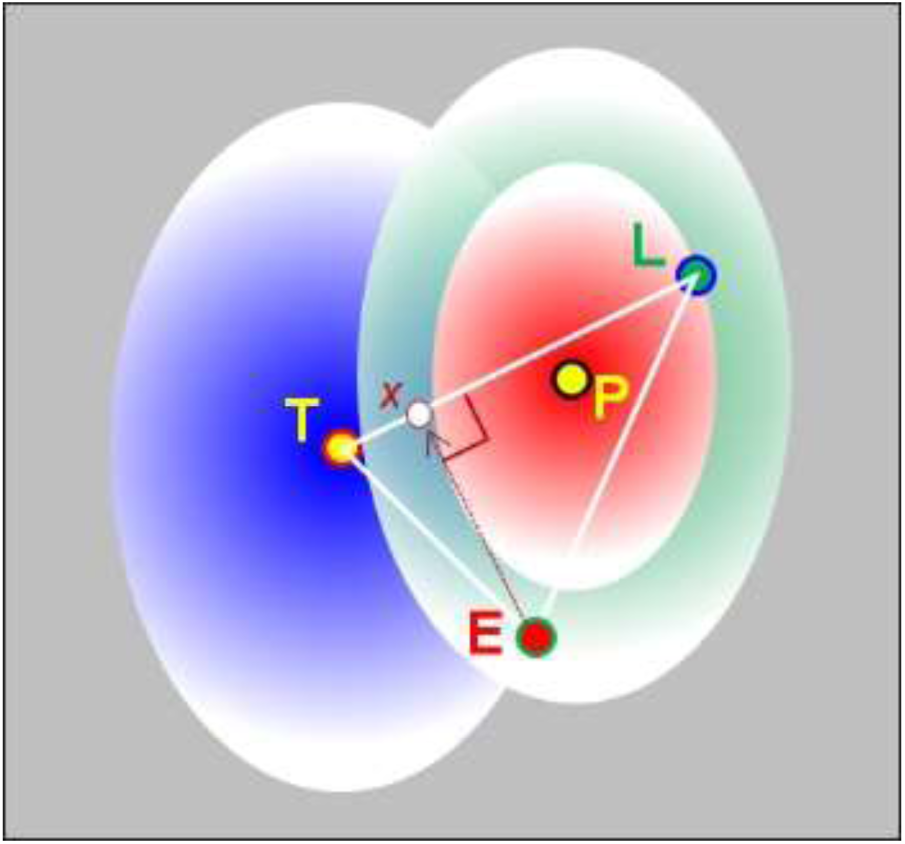
*A simple Internal Model. The blue, red, and green blurred ellipses represent the planning (P around the target Tlocation), landing (L around the P location), and the Estimated landing (E is also around P) uncertainties (with combined variances* 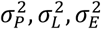 *see the Simulations section) as instantiated on each trial together with the corresponding LT̅, LE̅, and TE̅ norms (white triangle) and with the projection x of E on LT̅. The ellipses’ vertical axes correspond to the direction of the target relative to the starting point, which is also the direction of the hand movement.*

The variances of the three hypothesized distributions (σ_P_, σ_L_ σ_E_) can be obtained by solving the above set of equations. For the two-dimensional case, the set of equations can be duplicated, one set for each dimension. Given the constant shape found for the three involved distributions (LT, EL, and ET; see the Results), the two sets differ only by a scaling factor (with σ1_ij_ = r σ2_ij_); thus, their solutions are expected to differ by the same factor. Here, for simplicity, we present variances corresponding to the sum of variances over the two dimensions. The noise correlation between 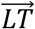 and 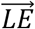 is predicted by their shared variance.

Assuming that the observed L and E coordinates are normally distributed (98% of pointing and estimation distribution passed the Kolmogorov-Smirnov test of normality at a level of p > 0.05), they are fully described by their mean and SD. If the model’s underlying assumptions are correct, the data (excluding the consistent errors) can be reconstructed with an accuracy limited only by the sampling rate (the number of trials). The outcome of model simulations using the derived model’s precision parameters (σ_P_, σ_L_ σ_E_) is presented against the empirical data in the Simulation section (Figure 8). Simulated and empirical data are in nearly perfect agreement, hence confirming the validity of the model’s assumptions. Note that Ol as well as Oe include some memory noise, with the first being more of the iconic memory type (because movement planning may have not been completed by the time of the target’s offset, i.e., 100 ms (Shadmehr et al., 2010) and with the latter of the short-term memory type.

**Figure 8.**
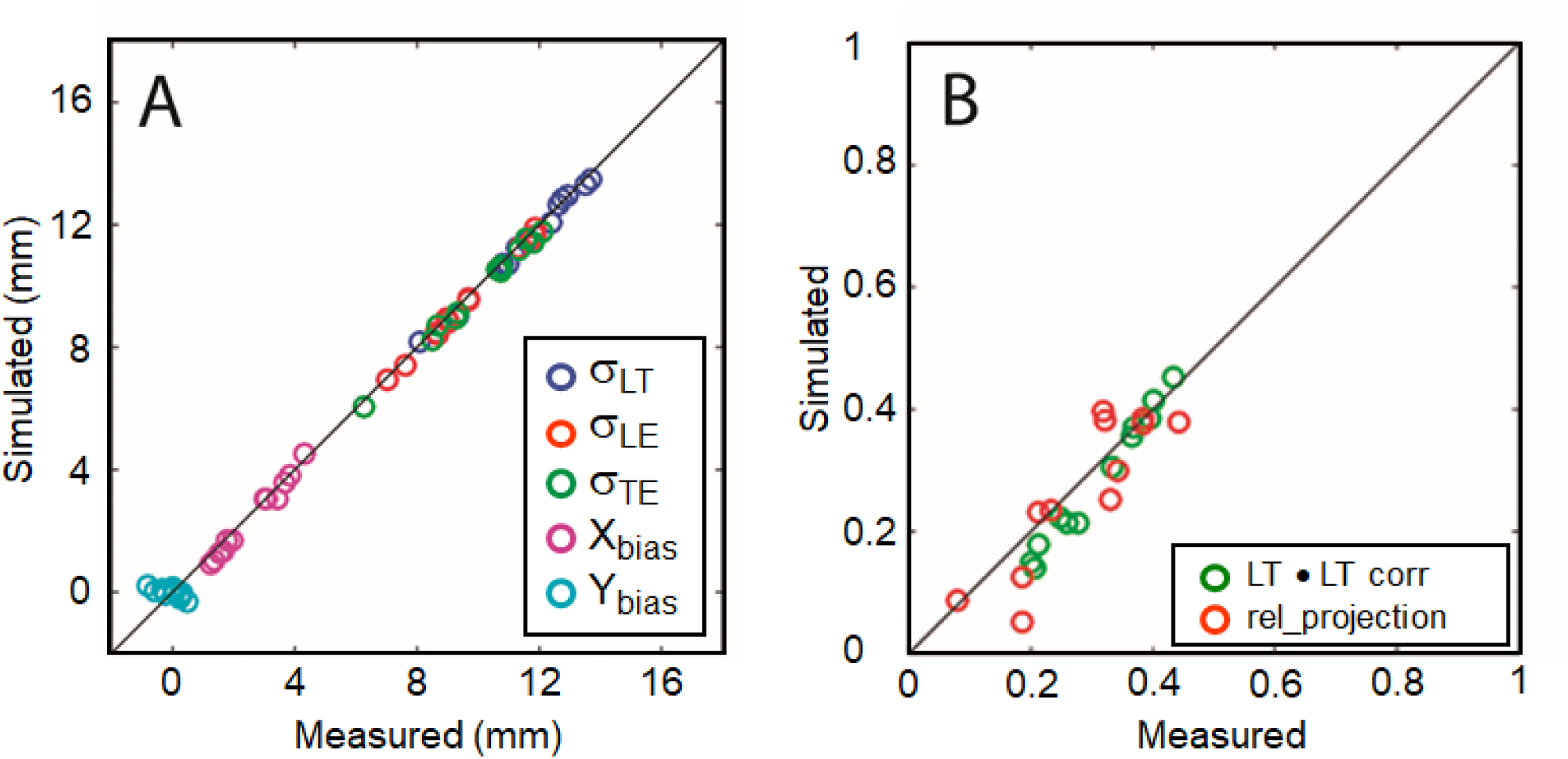
*A comparison between measured and simulated data. **A.** σ_LT_,σ_EL_ and σ_ET_, and of the E bias from L toward T. Linear correlations r > 0.99 for all sets but for the Ybias (r=0.47). **B.** Measured* vs. *simulated LT-LE vector correlations and relative projections (see Figure 5 in main text), with linear correlations of 0.99 and 0.88, respectively.*

Figure 7 presents these model parameters corresponding to the four experimental conditions and to the three T-arrays (marked as −1,0,1 for left, center, and right T-arrays) computed from Eq. 1 using the measured values of σ_LT_σ_EL_ and σ_ET_ averaged over all participants. The derived model’s parameters are intended to remap our directly measured random variables (characterized byσ_LT_σ_EL_ and σ_ET_ onto the hidden internal processes (characterized by σ_P_σ_L_ and σ_E_ note their symmetry with respect to the center T-array with larger SDs for the side T-arrays averaged across conditions by ∼35%). Target precision affects mainly σ_P_, which is ∼24% larger for the P-conditions than for the P+ conditions (two-tailed t-test, p=0.005), and to a smaller degree, σ_E_ which is increased by 8% under the P-condition (p=0.001), while having no appreciable effect on σ_L_ (∼1%, p=0.58). Vision of the hand during pointing has a strong effect on σ_L_, reducing it by ∼43%, as expected from studies having demonstrated online motor adjustments even for rapid pointing (Desmurget & Grafton, 2000; Gordon & Ghez, 1987; Paillard, 1996; Prablanc & Martin, 1992; Sabes, 2000; Sarlegna et al., 2004; Sarlegna & Sainburg, 2009; Saunders & Knill, 2003, 2004). Planning (1/σ_P_) and estimation (1σ_L_) precisions are expected to depend less on seeing the moving hand(Sarlegna et al., 2003, 2004). Indeed, both show small but significant gains from vision of respectively, ∼7% (p < 0.05, one-tailed pairwise t-test), and ∼13% (p<0.01, one-tailed pairwise t-test).

**Figure 7.**
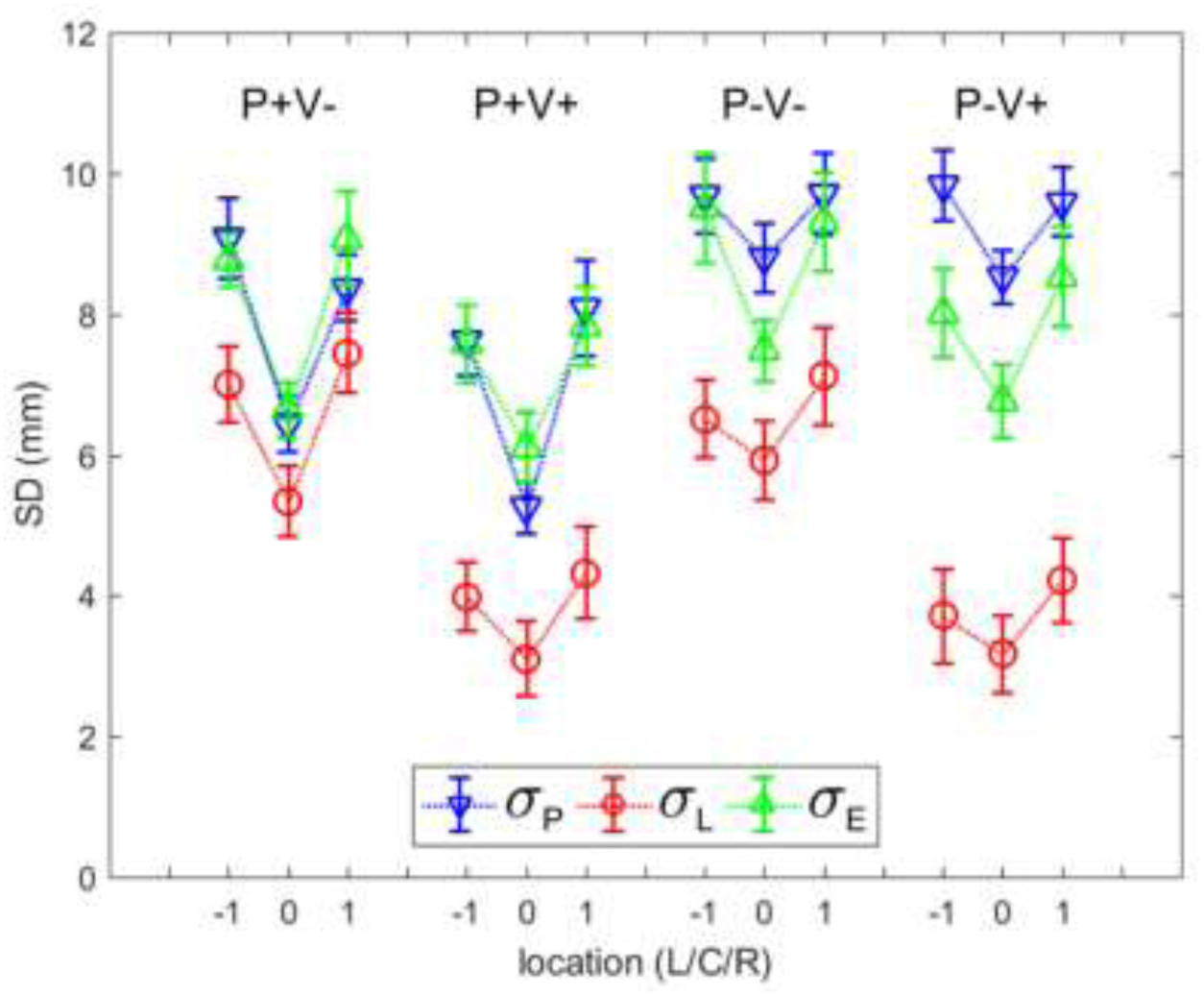
*The three standard deviations of the simple internal model (in mm), namely, of the planned landing (σ_P_), of the actual landing (σ_L_ and of the estimated landing ( σ_E_ inverted triangles, circles and upright triangles, respectively), as derived from the data averaged over all participants (Fig. 4) and used to simulate these data (see the Simulation section), for each of the four experimental conditions and for each of the 3 T-arrays (marked −1, 0, 1 for Left, Center, and Right; L/C/R). Error bars are ±1SEM (N=13).*

The model does not explicitly consider the contribution of vision to the planning and estimation precisions. However, its use of the measured L and E standard deviations (regarding T and L, respectively; eq. 1) as coefficients, generates visual gains. By comparison, the measured L (relative to T) and E (relative to L) gains due to vision are 17% and 22% (Fig. 4). According to the model (eq. 1), both of these visual gains are mostly due to an increased landing precision relative to P (σ_L_). Thus, independently of the fact that V+ conditions allow participants to see their landing location, the increase in E precision with vision is, according to the model, a direct consequence of an increase in L precision (σ_EL_^2^ = σ_E_^2^+σ_L_^2^)

## SIMULATIONS

A trial-by-trial simulation of our model (Fig. 6; eq. 1) was carried out for each experimental condition and for each of the three target arrays, that is 4 × 3 simulations. First, we found for each participant the model parameters (σ_P_, σ_L_ σ_E_ for each of the 12 simulations by solving Eq. 1 and using the empirical measurements of σ_LT_,σ_EL_ and σ_ET_ Next, these parameters, averaged across participants (Fig. 7), were used to generate artificial L and E locations for the T locations used in the experiments (140 locations × 13 participants = 1,820 trials/simulation/condition/target array). In the simulations, for each T location, a random generator (normal distribution, mean = 0, SD = σ_P_) created a P location around which the L and E locations were generated using independent random draws (normal distributions, means = 0, SD = σ_L_,σ_E_).In line with the observed results, showing non-isotropic distributions with equal shape for landing and estimation (see Results/Raw-data, and Model sections), all simulated normal distributions were assumed to be two dimensional with a ratio of 1.5 between their two standard deviations (so that the combined SD equals the derived model parameters). This is acceptable as the ellipses corresponding to landing (TL) and estimation (LE) had approximately the same orientations (for the 3 locations: 150, 94, 33 degrees vs 147, 92, 35 degrees). Furthermore, estimation distributions relative to target (TE) also showed equivalent orientations (144, 92, 31 degrees). Ellipses’ shapes also varied only marginally across distributions (differences in major to minor axes ratio <2%). This means that eq 1 (in the main text) can be written as two independent sets of equations describing the variances along the major and minor axes, with the two sets differing by only a scaling factor applied to the left hand side (leading to solutions that differ by the same scaling factor). The reported variances are obtained by adding the corresponding solutions of the two sets of equations.

The generated location triplets (T, L and E) were fed into the data analysis program to compute the simulated results. The comparison between measured and simulated results are presented in Figure 8A for σ_LT_,σ_EL_ and σ_ET_. Fig. 8A also presents a comparison between the measured and simulated E biases. These biases correspond to E being, on the average, away from L toward T (see Fig. 1), with the bias on the 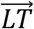 vector (X_bias_) and orthogonal to it (Y_bias_). Fig. 8B presents a comparison between two other measured properties of the data, presented in Fig. 5: the 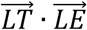 vector correlation, and the relative projection (X_bias_ relative to the 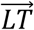 length, averaged across trials).

## DISCUSSION

The present study addressed the question of whether and how well one knows the outcome of one’s actions and, more specifically, of a speeded up pointing task. As indicated in the Introduction, this question pertains to people’s *cognitive apprehension* of the consequences of their motor actions rather than to their capacity for reproducing those actions. Fourneret & Jeannerod (1998) were the first to address this question. In their study, participants were requested to draw a visible continuous line on a tablet within less than 5 s in order to join a 22 cm distant target in the sagittal plane without seeing their hand. Unbeknown to them, a linear directional bias of varying direction was added to the visible line. Participants successfully reached the target by applying a deviation symmetrical to the added bias but failed to acknowledge that their hand was following this oppositely biased direction and asserted instead that it remained in the sagittal plane. The authors concluded that the participants were “poorly, if at all, aware of the details of their motor performance and (…) unable to correctly monitor, consciously, the signals generated by their own movements” (p 1137). Although this is tantamount to concluding that, had the participants not been allowed to see the final outcome of their action, they would have been unaware of it, Fourneret & Jeannerod’s experimental paradigm was not aimed at testing this assertion directly and as such, it did not allow this specific introspective knowledge to be fully characterized. In addition, whereas Fourneret & Jeannerod’s results strongly suggest that their participants’ introspective judgment relied entirely on their originally planned limb movement, their paradigm does not allow the putative discrimination between the actual end-point of the motor planning and the visible target they were meant to reach. This is precisely one of the achievements of the present paradigm and of the proposed modeling of our data. Fourneret & Jeannerod’s inferences have been recently modulated by a study that showed well above chance performances when the participants had to decide whether their landing location was left or right of the target (even without visual feedback), but, in accordance with the present results, they failed to appropriately correct their errors after completing the movement, a situation similar to the present estimation behavior (van Dam & Ernst, 2013).

The present results reveal some unexpected properties of the participants’ ability to estimate their own pointing action. First, the data reveal significant consistent errors in both pointing 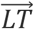 and estimation 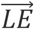. In particular, the finding that the landing estimation was consistently (i.e., across all experimental conditions) though idiosyncratically shifted away from the landing point suggests that participants have no or limited knowledge of their landing points. Surprisingly, despite these idiosyncratic, consistent errors, our noise analysis revealed a participant-independent behavior, namely, a significant bias of the estimated landing, E, away from the actual landing, L, and toward the target, T.

Our data show practically equal L (with respect to T) and E (with respect to L) errors with the Bayesian implication of quasi-flat likelihoods. Within the Bayesian framework, the L and E distributions would be referred to as prior and posterior, respectively. Given their observed quasi-equality (with 67 out of 156 cases – 13 participants × 4 conditions × 3 locations – i.e., 42.9%, where the posterior precision was poorer than that of the prior, and with at least 4 such cases out of 12 per participant), the Bayesian account of the data would predict that the mean of the posterior coincides with the mean of the prior, i.e., a 100% E shift toward T. However, this is definitely not the presently observed result.

It could be argued that the discrepancy of the Bayesian account of the present data is due to it not including the putative memory loss during the time lapse between landing and estimation, particularly given that our design allowed for a free estimation time. Note, however, that the alternative to this design would have been to impose on participants’ estimation time the same time constraints as in the speeded up landing task, i.e., to force them into a ‘ballistic’ pointing mode. This would have been a valid design had our question focused on the participants accuracy in repeating their original (reflexive/ballistic) motor actions. Instead, our question focused on the *cognitive apprehension of the outcomes of those motor actions.* In addition, note that no such memory loss was considered by Wolpe et al.(Wolpe et al., 2014) when they fit their data with a Bayesian model. Had such a memory loss been a factor that explained why the presently observed estimation noise was equal to and often larger than the landing noise, it should have also appeared in Wolpe et al.’s data (i.e., an estimation noise larger than the performance noise, which is just the opposite of their findings) and it would have played against their Bayesian model fit of their data. Since this was not the case, one can conclude that the memory noise in both their and the present study did not play a critical rσ_LE_. It has been indeed shown that spatial localization thresholds for a test line increase negligibly for intervals larger than about 200 ms between the offset of a fixation line and the display of the test line (White, Levi, & Aitsebaomo, 1992). (Our participants’ average estimation time was about 1 s.) Also note that differences between the present results and those of Wolpe et al.’s could be due to critical differences between the two tasks: Participants’ task in Wolpe et al.’s study was not to estimate their own pointing location, but rather, it involved participants’ interaction with a computer screen via a mouse to control a visual pointer. Moreover, Wolpe et al.’s task, unlike ours, involved a fixed target that participants were encouraged to fixate on, and that remained on the screen 250 ms *after* pointing. It is possible that under such conditions participants were biased toward the target, potentially in agreement with the Bayesian framework but not necessarily so as similar results could have been obtained by using some simple idiosyncratic rules.

For now, our theoretical approach, based on a quantitative noise analysis, posits that participants do not know the outcome of their action (unless they actually see it), but rather, when asked to introspect, refer to the outcome expected from their *planned* action as if they had cognitively simulated the original motor plan (Sheahan, Franklin, & Wolpert, 2016; Wolpert & Ghahramani, 2000; Wolpert, Ghahramani, & Jordan, 1995). The end product of both action (landing) and introspection (estimation) is the planned outcome of the original action perturbed respectively by motor execution noise and by simulation noise together with, most likely, some memory loss.

### Modeling precision and E bias

The proposed model is parameter-free and is based on three postulates: (1) participants’ pointing is not directed to the target T per se but to their planned landing location, P, a random variable centered on T with a variance contributed to by the visual localization of T and by the motor planning and memory noise; (2) as a consequence, the landing location, L, is a random variable centered on P (rather than on T) with a variance contributed to by the motor execution noise; (3) participants have limited knowledge of their landing location proper, L, so that their L-estimate, E, is also centered on P (a form of “exaggerated expectation of success” (Wolpe et al., 2014). This feature of the model agrees with previous proposals relating awareness of action to motor control processes, mainly motivated by observed abnormalities of motor awareness in clinical cases (Blakemore, Wolpert, & Frith, 2002; Frith, Blakemore, & Wolpert, 2000; Moore & Haggard, 2008).

The model is described by three equations relating three unknowns (the standard deviations of P, L, and E) to the measured standard deviations of L relative to T, of E relative to L and of E relative to T for each of the 4 conditions and for each of the 3 target locations. By virtue of the statistical properties of P, L, and E, the model generates the statistical properties of the empirical data, namely, the variances of the 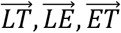 vectors, and the 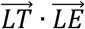 vector correlations across trials associated with the observed E bias away from L and toward T. As shown in the Simulation section, these model-derived features provide an excellent fit of those derived from the empirical data.

The isolated P, L, and E standard deviations computed by the model can be used to inspect the dependency of the related internal processes on the experimental conditions. We showed that the main effect of visual feedback lies in reducing the landing (L) variability. The derived average variability of the model’s σ_P_ variable for small targets in the presence of visual feedback (P+V+ condition) of about 8 mm - which was only slightly affected by visual feedback (see the Results) –, is quasi-identical to humans’ visual localization error of targets placed at 21 deg eccentricity in the absence of any contextual cues (e.g., screen edges) (White et al., 1992). For larger targets having increased spatial uncertainty, the corresponding σ_P_ is increased, as expected. Inasmuch as P is supposed to include planning noise, its variability (σ_P_) is expected to exceed the pure visual localization one. However, in contrast with Ref. 34, contextual cues were only partially obscured in the present experiments and the targets were presented at only 17 deg eccentricity; both differences are expected to decrease the P error. The estimation (E) and variability σ_L_ derived by the model is about the same under all experimental conditions, matching that of the P variable with a small target size (P+), indicating a limit set by visual acuity. In support of our basic conjecture that participants’ introspective knowledge reflects that of their planned rather than their actual landing locations, Collins, Rolfs, Deubel, & Cavanagh (2009) have shown that the displacement of a saccade target during the saccade is accurately judged independently of the saccade’s actual landing site, as if the observers’ spatial gauge was their planned saccade, i.e., less noisy than its execution.

Although the present model is descriptive, does not suggest a specific decision strategy, and is not intended to compete with the normative Bayesian framework, it suggests however that participants’ estimation of their landing location, which can be regarded as the posterior distribution, relies on their planned landing location, which in turn, can be regarded as a prior distribution. Within such a framework, vision of the pointing hand can be seen as contributing to shrinking the prior distribution and hence, decreasing the spread of the posterior as well as the estimation bias.

Certainly, the present results could still be modeled within the normative Bayesian framework but at the cost of yet another free parameter (in addition to the unknown prior) that would characterize the memory loss due to the free estimation time in our design. This would enable the Bayesian posterior to be smaller than the prior but would be at odds with Wolpe et al.’s finding that their Bayesian approach makes do without considering such memory noise (see above). Yet another alternative within the framework of our simplistic, parameter-free model is to assume that the motor plan is the prior and that the sensory information is not introspectively accessible. Positing such a limitation on sensory information is however contrary to the very founding principle of Signal Detection Theory (Green & Swets, 1966) and of the Bayesian framework (Ma, 2012; Ma & Jazayeri, 2014).

### Other features of the data

The data reveal other features of participants’ pointing and estimation behavior not intended to be addressed by our model. One is that the L and E precisions (respectively, around T and L) are correlated *across* participants: participants who are precise pointers are also precise estimators. This behavioral property, also reported for a task involving the timing of a button-press and its subsequent estimation, was accounted for by positing that participants scale their priors according to their motor precisions (Wolpe et al., 2014). Since the present data cannot be accounted for by a straightforward Bayesian model (i.e., without including an additional free parameter), and since the L and E precisions are independent random variables in the presently proposed model, the only reasonable interpretation of this result is that participants tend to match the precision of their estimation of their landing location with the precision of their sensory and motor execution capacities and that, as a consequence, they have implicit or explicit access to these precisions. Within the framework of the proposed model, whose fitted E and P precisions (respectively around P and T) reveal their quasi-equality, one can infer that participants do their best to match their landing estimation precision with that of their *motor planning* (rather than their sensory and motor execution), again with the implication that they have introspective access to this planning.

Another feature of the data which cannot be accounted for by the model, is the idiosyncrasy of the consistent L and E errors. The L consistent error may reflect participant-specific muscle constraints and/or systematic errors in the sensorimotor transformation from the visual representation of the target location to the joint-based motor command (Chang & Snyder, 2010; Flanders, Helms, & Soechting, 1992; Ghahramani, Wolpert, & Jordan, 1996; Sober & Sabes, 2003; Soechting & Flanders, 1989a, 1989b). Such systematic errors in the sensorimotor transformation should differentially affect the planning and the execution of the movement, on the one hand, and the estimation of the action outcome on the other, due, among other factors, to their timing differences (speeded up vs. time unconstrained behaviors, respectively).

## CONCLUSION

The present study assessed the precision and accuracy of participants’ knowledge of the outcome of one of their most frequent motor behaviors, namely, pointing to a visual target. The data and their modeling have established that this knowledge cannot be accounted for within a straightforward Bayesian framework and have shown instead that it can be fully accountable when positing that it is confounded with participants’ knowledge of their planned motor actions rather than with knowledge of the actual outcome of their actions. When asked to estimate their landing location, the participants seem to reenact their original motor plan.

## Acknowledgments

This research was supported by *The France-Israel Laboratory of Neuroscience* (to AG and DS), and by the Weizmann Braginsky Center for the Interface between the Sciences and the Humanities (to DS). The experiments were conducted by Esther Bernager, an undergraduate student under the supervision of A. Gorea.

